# Optimized implementations of voxel-wise degree centrality and local functional connectivity density mapping in AFNI

**DOI:** 10.1101/067702

**Authors:** R. Cameron Craddock, Daniel J. Clark

## Abstract

Degree centrality (DC) and local functional connectivity density (lFCD) are statistics calculated from brain connectivity graphs that measure how important a brain region is to the graph. DC (a.k.a. global functional connectivity density) is calculated as the number of connections a region has with the rest of the brain (binary DC), or the sum of weights for those connections (weighted DC). lFCD was developed to be a surrogate measure of DC that is faster to calculate by restricting its computation to regions that are spatially adjacent. Although both of these measures are popular for investigating inter-individual variation in brain connectivity, efficient neuroimaging tools for computing them are scarce. The goal of this Brainhack project was to contribute optimized implementations of these algorithms to the widely used, open source, AFNI software package.

## 1 Introduction

Degree centrality (DC) [1] and local functional connectivity density (lFCD) [2] are statistics calculated from brain connectivity graphs that measure how important a brain region is to the graph. DC (a.k.a. global functional connectivity density [2]) is calculated as the number of connections a region has with the rest of the brain (binary DC), or the sum of weights for those connections (weighted DC) [1]. lFCD was developed to be a surrogate measure of DC that is faster to calculate by restricting its computation to regions that are spatially adjacent [2]. Although both of these measures are popular for investigating inter-individual variation in brain connectivity, efficient neuroimaging tools for computing them are scarce. The goal of this Brainhack project was to contribute optimized implementations of these algorithms to the widely used, open source, AFNI software package [3].

## 2 Approach

Tools for calculating DC (`3dDegreeCentrality`) and lFCD (`3dLFCD`) were implemented by modifying the C source code of AFNI’s `3dAutoTcorrelate` tool. `3dAutoTcorrelate` calculates the voxel × voxel correlation matrix for a dataset and includes most of the functionality we require, including support for OpenMP [4] multithreading to improve calculation time, the ability to restrict the calculation using a user-supplied or auto-calculated mask, and support for both Pearson’s and Spearman correlation.

### 3dDegreeCentrality

Calculating DC is straight forward and is quick when a correlation threshold or is used. In this scenario, each of the .5*N_vox_*(*N*_*vox*_ — 1) unique correlations are calculated, and if they exceed a user specified threshold (default threshold = 0.0) the binary and weighted DC value for each of the voxels involved in the calculation are incremented. The procedure is more tricky if sparsity thresholding is used, where the top *P*% of connections are included in the calculation. This requires that a large number of the connections be retained and ranked - consuming substantial memory and computation. We optimize this procedure with a histogram and adaptive thresholding. If a correlation exceeds threshold it is added to a 50- bin histogram (array of linked lists). If it is determined that the lowest bin of the histogram is not needed to meet the sparsity goal, the threshold is increased by the bin-width and the bin is discarded. Once all of the correlations have been calculated, the histogram is traversed from high to low, incorporating connections into binary and weighted DC until a bin is encountered that would push the number of retained connections over the desired sparsity. This bin’s values are sorted into a 100-bin histogram that is likewise traversed until the sparsity threshold is met or exceeded. The number of bins in the histograms effects the computation time and determine the precision with which ties between voxel values are broken. A greater number of bins allow the sparsity threshold to be determined more precisely but will take longer to converge. Fewer bins will result in faster computation but will increase the tendency of the algorithm to return more voxels than requested. The chosen parameters enable ties to be broken with a precision of 1.0/(50 * 100), which in our experience offers quick convergence and a good approximation of the desired sparsity.

### 3dLFCD

lFCD was calculating using a region growing algorithm in which face-, side-, and corner-touching voxels are iteratively added to the cluster if their correlation with the target voxel exceeds a threshold (default threshold = 0.0). Although lFCD was originally defined as the number of voxels locally connected to the target, we also included a weighted version.

### Validation

Outputs from the newly developed tools were benchmarked to Python implementations of these measures from the Configurable Pipeline for the Analysis of Connectomes (C-PAC) [5] using in the publically shared Intrinsic Brain Activity Test-Retest (IBATRT) dataset from the Consortium for Reliability and Reproduciblity[6].

**Figure 1.**
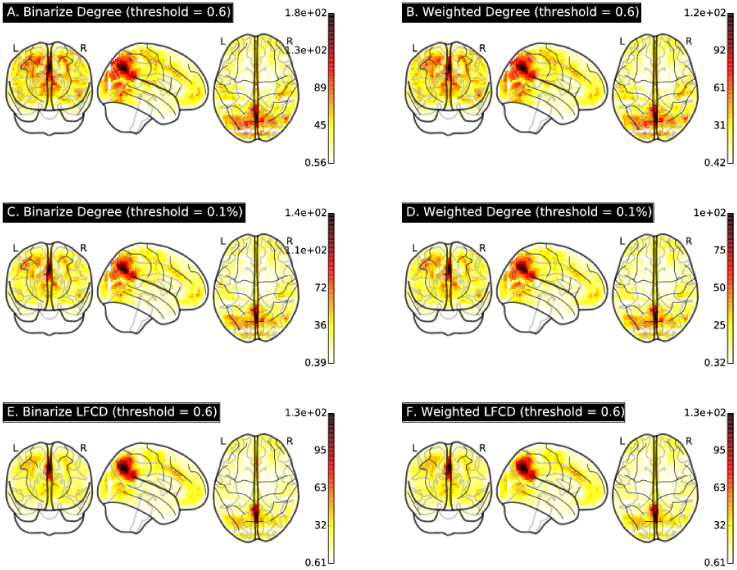
Whole brain maps of binarized and weighted degree centrality calculated with a correlation threshold of p > 0.6 (a-b) and sparsity threshold of 0.1% (c-d) and binarized and weighted lFCD calculated with a correlation threshold of p > 0.6 (e-f) averaged across maps calculated the first resting state scan of the first scanning session for all 36 participants’ data from the IBATRT data.

## 3 Results

AFNI tools were developed for calculating lFCD and DC from functional neuroimaging data and have been submitted for inclusion into AFNI. LFCD and DC maps from the test dataset (illustrated in Fig. 1) are highly similar to those calculated using C-PAC (spatial concordance correlation [7] p > 0.99) but required substantially less time and memory (see Table 1).

**Table 1.**
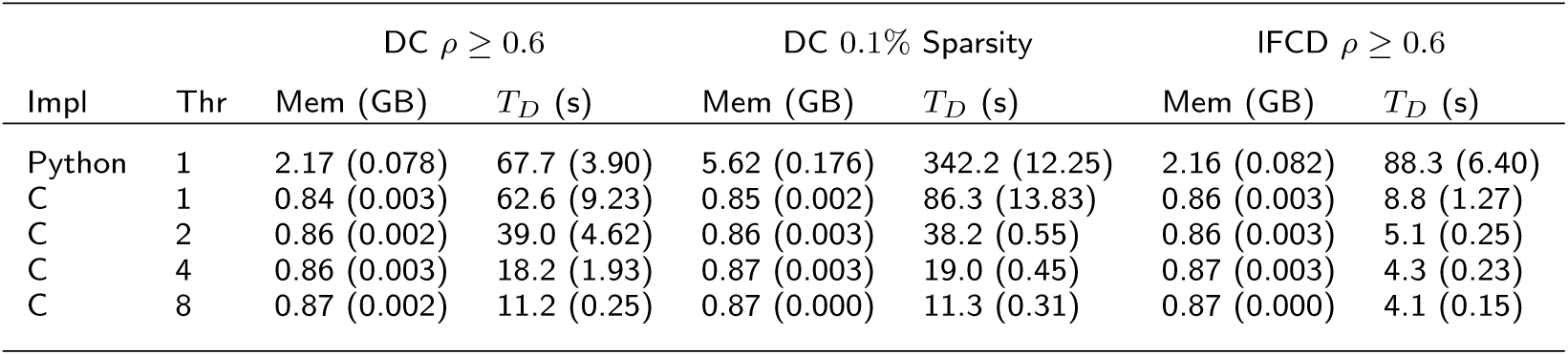
Comparison of the time and memory required by the Python and C implementations to calculate DC (sparsity and correlation threshold) and lFCD on the first resting state scan of the first scanning session for all 36 participants’ data in the IBATRT dataset. Values are averaged across the 36 datasets and presented along with standard deviations in parenthesis. Impl: Implementation, Thr: Number of threads used to process a single dataset, Mem: average (standard deviation) memory in gigabytes used to process a single dataset, *T_D_*: the average (standard deviation) time in seconds to process a dataset. These statistics were collected on a C3.xlarge Amazon Web Services Elastic Compute Cloud node with 8 hyperthreads and 15 GB of RAM.

| Impl | Thr | DC $\rho \geq 0.6$ | | DC 0.1% Sparsity | | IFCD $\rho \geq 0.6$ | |
| --- | --- | --- | --- | --- | --- | --- | --- |
| | | Mem (GB) | $T_D$ (s) | Mem (GB) | $T_D$ (s) | Mem (GB) | $T_D$ (s) |
| Python | 1 | 2.17 (0.078) | 67.7 (3.90) | 5.62 (0.176) | 342.2 (12.25) | 2.16 (0.082) | 88.3 (6.40) |
| C | 1 | 0.84 (0.003) | 62.6 (9.23) | 0.85 (0.002) | 86.3 (13.83) | 0.86 (0.003) | 8.8 (1.27) |
| C | 2 | 0.86 (0.002) | 39.0 (4.62) | 0.86 (0.003) | 38.2 (0.55) | 0.86 (0.003) | 5.1 (0.25) |
| C | 4 | 0.86 (0.003) | 18.2 (1.93) | 0.87 (0.003) | 19.0 (0.45) | 0.87 (0.003) | 4.3 (0.23) |
| C | 8 | 0.87 (0.002) | 11.2 (0.25) | 0.87 (0.000) | 11.3 (0.31) | 0.87 (0.000) | 4.1 (0.15) |

## 4 Conclusions

Optimized versions of lFCD and DC achieved 4× to 10× decreases in computation time compared to C-PAC’s Python implementation and decreased the memory footprint to less than 1 gigabyte. These improvements will dramatically increase the size of Connectomes analyses that can be performed using conventional workstations. Making this implementation available through AFNI ensures that it will be available to a wide range of neuroimaging researchers who do not have the wherewithal to implement these algorithms themselves.

## Availability of Supporting Data

More information about this project can be found at: http://github.com/ccraddock/afni. Further data and files supporting this project are hosted in the *GigaScience* repository doi:10.5524/100219.

## Author’s contributions

RCC and DJC wrote the software, DJC performed tests, and DJC and RCC wrote the report.

## Acknowledgements

The authors would like to thank the organizers and attendees of Brainhack MX and the developers of AFNI. This project was funded in part by a Educational Research Grant from Amazon Web Services.

